# Reward Enhances Connectivity between the Ventral Striatum and the Default Mode Network

**DOI:** 10.1101/2021.07.28.454086

**Authors:** Ekaterina Dobryakova, David V. Smith

**Affiliations:** Center for TBI Research, Kessler Foundation, 120 Eagle Rock Ave, East Hanover, NJ, USA; Department of Psychology & Neuroscience, Temple University, Philadelphia, PA, USA

## Abstract

The default mode network (DMN) has been theorized to participate in a range of social, cognitive, and affective functions. Yet, previous accounts do not consider how the DMN contributes to other brain regions depending on psychological context, thus rendering our understanding of DMN function incomplete. We addressed this gap by applying a novel network-based psychophysiological interaction (nPPI) analysis to the reward task within the Human Connectome Project. We first focused on the task-evoked responses of the DMN and other networks involving the prefrontal cortex, including the executive control network (salience network) and the left and right frontoparietal networks. Consistent with a host of prior studies, the DMN exhibited a relative decrease in activation during the task, while the other networks exhibited a relative increase during the task. Next, we used nPPI analyses to assess whether these networks exhibit task-dependent changes in connectivity with other brain regions. Strikingly, we found that the experience of reward enhances task-dependent connectivity between the DMN and the ventral striatum, an effect that was specific to the DMN. Surprisingly, the strength of DMN-VS connectivity was correlated with personality characteristics relating to openness. Taken together, these results advance models of DMN by demonstrating how it contributes to other brain systems during task performance and how those contributions relate to individual differences.

## Introduction

A cardinal goal of neuroscience relates to characterizing how functional integration organizes discrete brain regions into cohesive networks that shape behavior (Park & Friston, 2013). Several distinct networks have been identified in both resting and task states (S. M. Smith, et al., 2009). Of these networks, the default mode network (DMN)—including the posterior medial cortex, medial prefrontal cortex, and lateral temporal-parietal regions—may be unique in its elevated baseline energy consumption (R. L. Buckner, Andrews-Hanna, & Schacter, 2008; Randy L Buckner & DiNicola, 2019; Raichle, et al., 2001). Numerous studies, particularly earlier ones, noted that activation of the DMN appears to be highest when participants are engaged in inward-directed thought and lowest when participants are engaged in externally-directed tasks requiring focused attention (Hasenkamp, Wilson-Mendenhall, Duncan, & Barsalou, 2012; Hayden, Smith, & Platt, 2009; Scheibner, Bogler, Gleich, Haynes, & Bermpohl, 2017). In addition, while DMN has been shown to be engaged in a number of task-related behaviors with increases in neural activity in response to these behaviors (Smallwood, et al., 2021), the degree of task-related deactivation has been linked to performance (Anticevic, Repovs, Shulman, & Barch, 2010; Vatansever, Menon, Manktelow, Sahakian, & Stamatakis, 2015), suggesting a key role for the DMN in shaping behavior. Despite these advances, we still know very little about how task-related changes in DMN are linked to changes in effective connectivity with other brain regions.

Some studies have begun to point to a key link between the DMN and the striatum. For example, one study using physio-physiological interaction analyses demonstrated that the striatum interacts with the salience network and the DMN (Di & Biswal, 2014). Other studies have shown that striatal dopamine levels are linked to the magnitude of DMN deactivation during visuospatial attention (Tomasi, et al., 2009) and tasks requiring cognitive flexibility (Dang, Donde, Madison, O’Neil, & Jagust, 2012). In addition, DMN responses have also been associated with gamma oscillations in the striatum, which may serve to facilitate switching between externally-driven and internally-driven brain states (Nair, et al., 2018). While these observations illustrate some indirect relationships between the DMN and the striatum, it remains unclear whether there are contributions from the DMN to the striatum during a task. Observing this type of effective connectivity would lend support to the emerging idea that a key function of the DMN resides with guiding goal-directed behavior and value-based decision making (Dohmatob, Dumas, & Bzdok, 2020).

To further investigate DMN-striatum connectivity in healthy individuals, we examined connectivity with the DMN during a reward processing task. Thus, our analyses focused on two key questions. First, does the receipt of reward drive connectivity between the DMN and striatum? We hypothesized that the psychological context of reward consumption would alter effective connectivity between the DMN and the striatum. Due to previous studies showing DMN to overlap with regions involved in value computation, as during reward processing (Acikalin, Gorgolewski, & Poldrack, 2017; Gusnard, Akbudak, Shulman, & Raichle, 2001), we expected to observe increased coupling between the DMN and the striatum. We used data from the Human Connectome Project and applied a network-based psychophysiological interaction (nPPI) analysis (Dominic S Fareri, Smith, & Delgado, 2020; Utevsky, Smith, Young, & Huettel, 2017). This analytical approach merges canonical psychophysiological interaction (PPI) analysis (K. J. Friston, et al., 1997; McLaren, Ries, Xu, & Johnson, 2012; D. V. Smith, Gseir, Speer, & Delgado, 2016) with dual-regression analysis (Filippini, et al., 2009; Nickerson, Smith, Ongur, & Beckmann, 2017; D. V. Smith, et al., 2014) to assess how networks interact with other regions as a function of task context.

Second, if there is reward-dependent connectivity between the DMN and striatum, is it associated with behavior and personality characteristics that are functionally relevant to reward processing? Many studies have investigated the function of DMN through the lens of individual differences and linked certain personality characteristics to brain regions related to dopaminergic circuits (Käckenmester, Bott, & Wacker, 2019; Passamonti, et al., 2015). Similarly, numerous studies have linked the DMN to specific personality factors (Cai, Zhu, & Yu, 2020; Markett, Montag, & Reuter, 2018; Nostro, et al., 2018) and to openness specifically (Roger E Beaty, et al., 2016; Marstrand-Joergensen, et al., 2021; Simon, Varangis, & Stern, 2020), a personality factor linked to sensation seeking and new experiences. We thus examined whether Neuroticism-Extraversion-Openness Inventory Five-Factor Inventory (NEO-FFI) is associated with reward-dependent connectivity strength with DMN. In addition, we examined the association between reward-dependent connectivity strength with DMN and delayed discounting since previous work showed that the ability to wait for a larger reward is positively associated with mind-wandering (Smallwood, Ruby, & Singer, 2013), a behavior that characterizes DMN.

## Methods

### Participants

We obtained behavioral and neuroimaging data of 495 randomly selected subjects from the Human Connectome Project (HCP; www.humanconnectome.org), an open-access database aimed at collecting healthy participant data from over 1,200 people. Although recent work has shown that effect sizes for reward-related striatal activation in the HCP is modest with a median d = 0.40 (Poldrack, et al., 2017), we anticipated effect sizes associated with connectivity would be much smaller. Yet, even with a smaller expected effect size, we reasoned that a subsample of at least 376 participants would be sufficient for detecting small effects (d = 0.20) with 90% power and a two-sided paired t-test with alpha = 0.01. While less is known about the effect sizes for individual differences in brain-behavior relationships, there is good reason to believe such effects are generally small. For example, a recent examination of 87 meta-analyses on individual differences revealed a median r = 0.19 (Gignac & Szodorai, 2016). Detecting this effect with 90% power and alpha = 0.01 would require at least 405 participants. Within our subsample of 495 participants, ages ranged from 22 to 35 years old (Barch, et al., 2013).

Because of technical difficulties and excessive head motion, 457 out of the 495 downloaded subjects were used in the final analysis (see below for details). Within this sample, familial relationships (e.g., twins, siblings, etc.) were unknown for 4 individuals [including one participant without NEO (Neuroticism, Extraversion, Openness) data]; these participants were excluded from our analysis since it was impossible to account for family structure (Winkler, Ridgway, Webster, Smith, & Nichols, 2014). After all exclusions, our final sample consisted of 453 participants (females = 270).

All participants provided written informed consent in accordance with local Institutional Review Board policies. In addition, we also received permission to analyze the data in accordance with HCP ethics principles.

### Behavioral Paradigm

Participants completed a card-guessing task adapted from previous work (Delgado, Nystrom, Fissell, Noll, & Fiez, 2000) where they had to guess whether the number on a facedown card was higher or lower than 5. Following their response, participants were presented with one of three feedback options: 1) a green up arrow indicating a correct guess resulting in a gain of $1.00; 2) a red down arrow indicating an incorrect guess resulting in a loss of $0.50; or 3) a gray double-headed arrow for neutral trials. Stimuli were presented in blocks of eight trials. Trial blocks were either mostly reward or mostly loss, including six of the main conditions and two randomized trials from the remaining conditions. Each task block lasted 28 seconds and was separated by fixation blocks lasting 15 seconds. Two runs were completed, composed of 2 mostly reward, 2 mostly loss, and 4 randomly placed fixation blocks. The facedown card with a “?” was presented for up to 1.5 sec, with 1 sec allotted for feedback and a 1 sec inter-trial interval for the presentation of a fixation cross. All participants received standardized monetary compensations after completing the task. See Barch et al., (2013) for more details.

### fMRI preprocessing

Our data preprocessing and analysis was carried out using FMRIB Software Library tools (v6.00; www.fmrib.ox.ac.uk/fsl). For the analyses in the current paper, we used the minimally preprocessed data, the end product of the volume-based fMRI pipeline, which were motion corrected and normalized to the MNI template (Glasser, et al., 2013). We excluded 31 subjects whose average relative volume-to-volume motion qualified as an outlier relative to other subject; we identified outlier subject using a standard boxplot threshold (i.e., the 75^th^ percentile plus 1.5*lnterquartile Range (IQ.R); IQR for right-to-left phase encoding scan: 0.154; IQR for left-to-right phase encoding scan: 0.174). We excluded an additional 7 subjects due to technical difficulties (e.g., corrupt downloads).

Further, we removed the first 15 volumes to achieve steady state magnetization, and applied spatial smoothing with a 4 mm full-width at half-maximum (FWHM) Gaussian kernel. The output of the preprocessing was further processed with ICA-AROMA (Pruim, et al., 2015), which removed motion components from the dataset. Although the details of the ICA-AROMA approach are contained in the original report (Pruim et al., 2015), we briefly reproduce the key details here. First, we conducted a probabilistic independent component analysis (ICA) (Beckmann & Smith, 2004) as implemented in MELODIC (Multivariate Exploratory Linear Decomposition into Independent Components) Version 3.10, part of the FSL software package, for each participant. Prior to estimating each ICA, the input data were demeaned and the variance was normalized across voxels. The number of dimensions (i.e., the number of components) was estimated using the Laplace approximation to the Bayesian evidence of the model order (Beckmann & Smith, 2004). The whitened observations were then decomposed into sets of vectors that describe signal variation across the temporal domain (time-courses) and across the spatial domain (maps) by optimizing for non-Gaussian spatial source distributions using a fixed-point iteration technique (Hyvarinen 1999). We then normalized the estimated component maps by dividing the maps by the standard deviation of the residual noise and threshold using a mixture-modeling approach that gives equal weight to false negatives and false positives (Beckmann & Smith, 2004). Normalized and thresholded component maps were then submitted to a classifier to automatically label components as signal and head motion (Pruim et al., 2015). As detailed in the original ICA-AROMA paper, this approach assumes that independent components labeled as motion will exhibit edge artifacts (i.e., ringing), shared variance with head motion parameters, and/or activation in white matter and cerebral spinal fluid. Finally, independent components labeled as motion were then filtered from each dataset using regression (Pruim et al., 2015). After ICA-AROMA, the denoised dataset was further temporally filtered (high-pass temporal filtering 90 sec) prior to statistical analysis.

### fMRI analysis

Statistical analysis was carried out using general linear models (GLMs) with pre-whitening with local autocorrelation correction (Woolrich, Ripley, Brady, & Smith, 2001). We specifically examined GLMs to assess activation and connectivity associated with the task.

Our GLM assessing activation consisted of two regressors corresponding to reward and punishment blocks (duration = 28 seconds). Each task regressor was convolved with a double-gamma representing the hemodynamic response function. Our primary contract vectors quantified the relative activation for reward compared to punishment.

We extended this GLM-based approach to assess psychophysiological interactions (PPI) linked to task-dependent connectivity (K. J. Friston, et al., 1997). We note that, unlike functional connectivity analyses—which could reflect changes in another connection, observational noise, or neuronal fluctuations (K. J. Friston, 2011)—PPI is a simple test for effective connectivity. Indeed, PPI analyses are based on an explicit linear model of coupling between one or more brain regions. As described in detail elsewhere (K. J. Friston, et al., 1997; D. V. Smith, et al., 2016), this PPI model allows researchers to test for directed changes in connectivity by establishing a significant interaction between the seed region (or network) and the psychological context (e.g., reward relative to punishment). Although the direction of the change is explicitly specified in the model, we acknowledge that the post hoc interpretation of the results can be ambiguous and can be further refined with dynamic causal modeling and then conducting Bayesian model comparisons (K. Friston, 2009; K. J. Friston, Harrison, & Penny, 2003).

We estimated four separate network PPI (nPPI) models. Our primary model was focused on task-dependent changes in connectivity with the DMN. In addition, we also estimated three additional nPPI models using networks linked to cognition and executive function: executive control network (ECN), left fronto-parietal network (lFPN), and right fronto-parietal network (rFPN). All networks were taken from prior work (S. M. Smith, et al., 2009). Each model used the generalized PPI model (McLaren, et al., 2012) with a total of 14 regressors: network time course, reward x network time course interaction, and punishment x network time course interaction, and the time courses of nine additional networks identified in prior work (Smith et al., 2009). Including the time courses of these additional networks controls for the influence of other networks and renders more accurate estimates of effective connectivity (K. J. Friston, 2011).

To average contrast estimates over runs within each subject, we carried out a fixed effects analysis by forcing the random effects variance to zero in FLAME (FMRIB’s Local Analysis of Mixed Effects) (Beckmann, Jenkinson, & Smith, 2003; Woolrich, Behrens, Beckmann, Jenkinson, & Smith, 2004). We assessed statistical significance of group results using PALM (permutation analysis of linear models), a nonparametric permutation-based cluster-extent thresholding method, with 5000 random permutations and with cluster-forming threshold set to z > 2.3 (Eklund, Nichols, & Knutsson, 2016; Winkler, et al., 2014). We specifically used PALM because it allowed us to control for family structure, account for the effects of heritability that would make data points from related individuals non-independent, and restricts the number of times data can be permuted.

### Network Identification and Extraction

To obtain an independent estimate of network connectivity during reward processing, we used the network maps from Smith et al. (2009). As detailed in the original paper (Smith et al., 2009), these maps were created using a spatial ICA of resting-state data, which were then matched via correlation to task-based networks that emerged from activation coordinates. These maps are continuous (I.e., they are not restricted to discrete regions) and are thus perfect inputs into dual regression analysis (Nickerson et al., 2017). To extract whole-brain time courses associated with each RSN, we ran spatial regression. Spatial regression was performed as part of dualregression analysis (e.g. (Filippini, et al., 2009; Gordon, Stollstorff, & Vaidya, 2012; D. V. Smith, Sip, & Delgado, 2015; D. V. Smith, et al., 2014). The first step of the dual-regression analysis regresses the functional data of each participant (input) onto a set of spatial maps (RSNs from prior work in our case; Figure 1). Each of these spatial regressions results in an estimated time course of activation for each network (output). In the second step of the dual regression, these time courses (input) are then used as the “physiological” regressors in a conventional PPI model (as described below), which allows us to test specific hypotheses about task-dependent connectivity with each network of interest, such as the DMN (output). We note that this process can alternatively be run with group-level ICA maps, rather than maps from another study. Although maps derived from a group-level ICA can offer additional denoising (Nickerson et al., 2017), our prior work has shown a great deal of similarity between results derived from the Smith et al. (2009) maps and a group-level ICA (Smith et al., 2014; Fareri et al., 2021).

**Figure 1:**
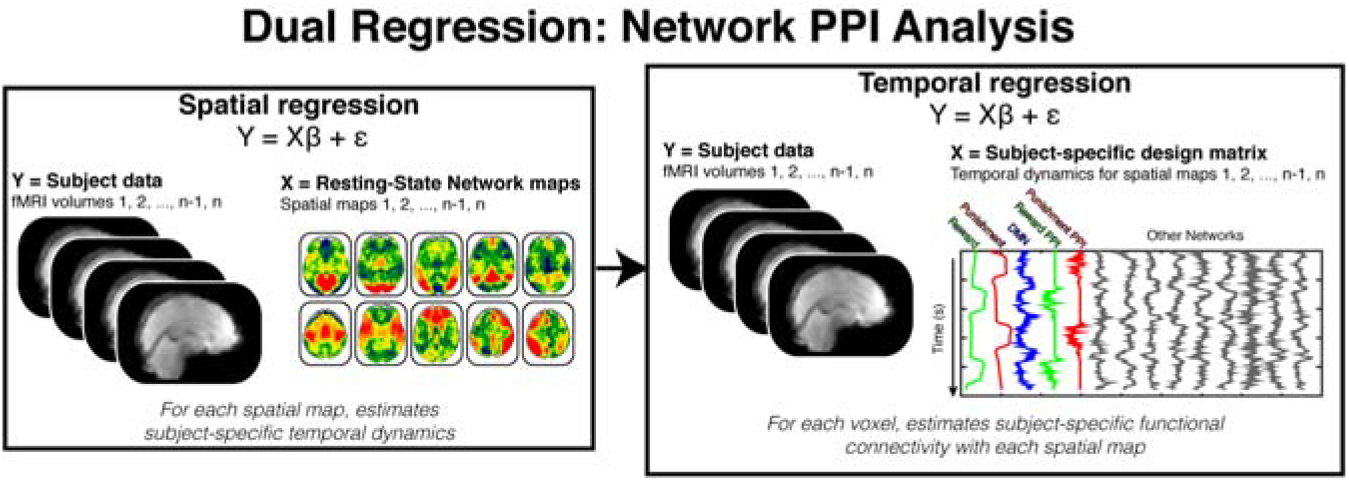
Analysis Schematic. Our network psychophysiological interaction analysis (nPPI) approach builds off of the dual-regression approach in that in consists of two key stages. In the first stage, the functional data from each participant is regressed onto a set of spatial maps reflecting common networks found in both resting-state and task-based data. This spatial regression results in a set of subject-specific temporal dynamics for each network. Note that the spatial maps shown here are derived from prior work (Smith et al., 2009). In the second stage, the temporal dynamics for each network are included in standard general linear model along with the task regressors (temporal regression). Regressors corresponding to psychophysiological interaction terms are created by multiplying each task regressor by the network of interest (i.e., default mode network).

In addition, we conducted a control seed-based PPI analysis to determine if a particular seed contributes to the observed DMN-VS connectivity. We used representative nodes of the DMN by identifying the peaks within the network map (Smith et al., 2009). These peaks were located in the ventromedial PFC (VMPFC), posterior cingulate cortex (PCC), and left and right temporo-parietal junction (lTPJ and rTPJ, respectively); and each peak served as a seed region in a conventional seed-based PPI analysis.

### Individual differences data

HCP provides rich behavioral data related to personality factors and cognition. However, we decided to focus on data related to impulsivity and personality as these constructs have been previously shown to be associated with DMN (e.g., (Cai, et al., 2020; Sampaio, Soares, Coutinho, Sousa, & Gonçalves, 2014a)). To examine whether DMN connectivity during reward processing would show an association with cognitive and personality constructs, we performed multiple regression analysis between behavioral measures obtained outside of the scanner and connectivity results. Specifically, we examined impulsivity as assessed by delayed discounting task and personality as assessed by Neuroticism/Extroversion/Openness Five Factor Inventory (NEO-FFI). We used area-under-the-curve discounting measure from the delayed discounting task as it provides a summary measure about the degree one discounts delayed rewards. From the NEO-FFI, a validated personality inventory (Costa & McCrae, 2008), we used subscales that assesses individual’s degree of agreeableness, openness, conscientiousness, neuroticism, and extraversion. While the regression analysis focused on NEO-FFI and delayed discounting, in addition to NEO-FFI, and discounting measures, the regression included mean relative motion, VS activation associated with positive vs. negative reward presentation to control for the effect of these parameters in the model. These independent variables were all entered together in one step with DMN-VS connectivity as dependent variable. The regression included mean relative motion, NEO-FFI (agreeableness, openness, conscientiousness, neuroticism, extraversion), and discounting measures.

## Results

### Reward-Related activation

As a manipulation check, we examined whether there is differential activation in the striatum during reward processing. Consistent with prior studies using the card task (e.g., (Delgado, et al., 2000; Tricomi, Delgado, McCandliss, McClelland, & Fiez, 2006), we observed that the receipt of reward in comparison to punishment, evoked greater activation in the striatum and other regions (p>0.001, z > 3.1, corrected for multiple comparisons; Table 1). This finding also furthers previously reported results from the HCP data (Barch, et al., 2013). No regions survived the threshold during the punishment vs. reward comparison.

**Table 1.** Brain regions showing greater activation to reward vs. punishment.

| Region | Number of Voxels | Hemisphere | Z-stat | Peak X | Peak Y | Peak Z |
| --- | --- | --- | --- | --- | --- | --- |
| Occipital Pole | 23088 | R | 16.1 | 14 | -96 | 18 |
| Ventral Striatum | 4560 | R | 9.58 | 6 | 14 | -2 |
| Dorsomedial Prefrontal Cortex | 2266 | L | 7.35 | -44 | 8 | 32 |
| Supramarginal gyrus | 157 | L | 4.66 | -50 | -48 | 16 |
| Amygdala/parahippocampal gyrus | 114 | L | 4.75 | -30 | -10 | -14 |
| Supplementary Motor Area | 107 | L | 5.1 | -6 | 6 | 56 |
| Posterior Lobe, Cerebellum | 83 | R | 6.14 | 20 | -42 | -46 |
| Paracingulate gyrus | 65 | L | 4.34 | -8 | 24 | 44 |
| Putamen | 48 | R | 4.96 | 28 | -12 | 10 |
| Parahippocampal Gyrus | 48 | L | 5.19 | -16 | -30 | -6 |
| Parahippocampal Gyrus | 39 | R | 4.94 | 8 | -32 | -4 |
| Middle Temporal Gyrus | 39 | R | 4.39 | 48 | -10 | -14 |
| Caudate Tail | 33 | R | 4.49 | 22 | -18 | 24 |
| Putamen | 32 | R | 4.77 | 32 | -10 | -10 |

We also examined how reward and punishment modulated responses of the DMN. To do this, we extracted the network responses using the spatial regression component of dual regression analysis (Dominic S Fareri, et al., 2020; Nickerson, et al., 2017; D. V. Smith, et al., 2015). To facilitate comparison with other studies, this spatial regression utilized unthresholded whole-brain spatial maps from prior work (Smith et al., 2009). Importantly, the resulting time courses (network responses) are derived without knowledge of our task structure and hence there can be no concern regarding circularity (Nickerson et al., 2017). Network responses were then regressed onto a model containing the predicted hemodynamic responses for reward and punishment blocks. Consistent with prior work, we found that the DMN was strongly deactivated during the task (Figure 2; t(453) = −75.31, p < 0.0001), but the magnitude of deactivation did not significantly differ between reward and punishment blocks (t(453) = 1.35, p = 0.1792). For completeness, we also assessed task-related responses in other networks that have been theorized to play and antagonist role with the DMN (Fox, et al., 2005; Uddin, Clare Kelly, Biswal, Xavier Castellanos, & Milham, 2009). These networks included the executive control network (ECN) and the left and right frontoparietal networks (lFPN and rFPN). Each of these networks exhibited a positive response to the task (ECN: t(453) = 2.37, p = 0.0184; lFPN: t(453) = 24.11, p < 0.0001; rFPN: t(453) = 37.57, p < 0.0001). Although the response to reward relative to punishment was larger in the ECN (t(453) = 3.52, p = 4.7838e-04) and the lFPN (t(453) = 4.71, p = 3.3716e-06), we found that the rFPN exhibited a larger response to punishment compared to reward (t(453) = −2.55, p = 0.0110). While some reward-related processes have been shown to be lateralized in some contexts (Palminteri, Boraud, Lafargue, Dubois, & Pessiglione, 2009), such differences are beyond the scope of our paper, as they could be driven by the single-handed button-press response procedure in the HCP.

**Figure 2.**
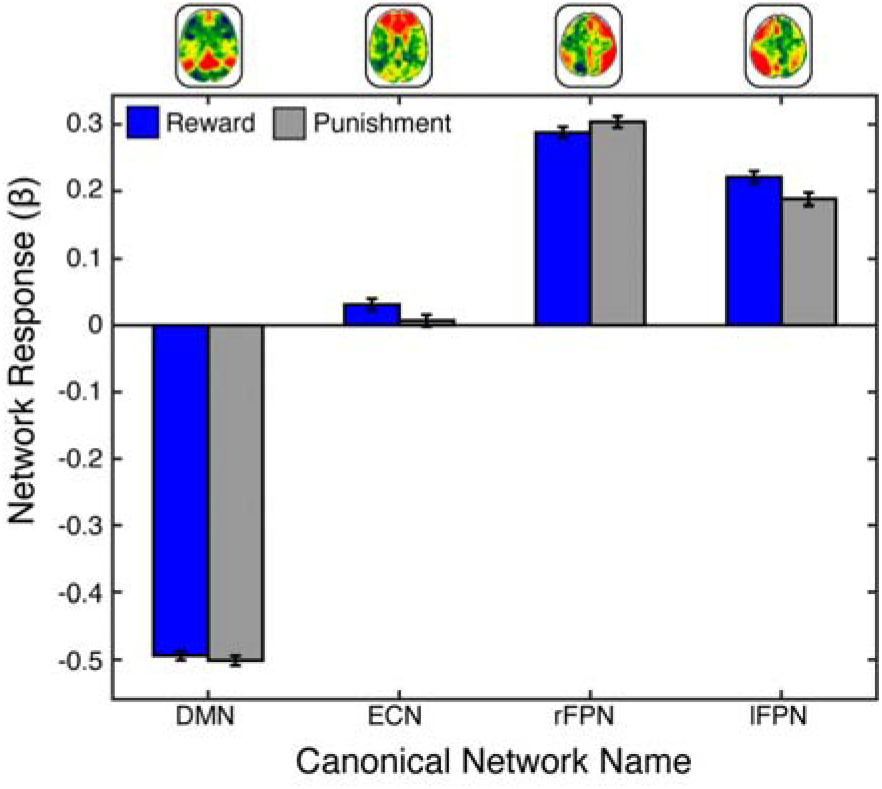
Network Responses to Reward and Punishment. We used spatial maps from prior work (Smith et al., 2009) to serve as networks in the current study (top: unthreshold maps with red depicting positive weighting and blue depicting negative weighting). Responses of these whole-brain maps were extracted using a spatial regression, which revealed the time course of each network within each participant in our sample. Next, we examined task-related responses for each network by regressing its activity onto a general linear model containing regressors for reward and punishment. We found that networks exhibited positive response to the task. As expected, we also found that the default mode network (DMN) was strongly deactivated during both reward and punishment. Abbreviations: DMN = default mode network; ECN = executive control network; rFPN = right frontoparietal network; lFPN = left fronto-parietal network.

### Reward Enhances Coupling Between the DMN and Ventral Striatum

Using the nPPI analysis, we examined reward-dependent connectivity of the DMN and the DMN showed increased connectivity with the left ventral striatum (VS) during the receipt of reward (*M* = 0.79, *S.E.M*. = 0.19) relative to punishment (*M* = −0.77, *S.E.M*. = 0.20) (Figure 3). Additionally, the DMN showed significant connectivity with the occipital pole and occipital fusiform gyrus. This result adds to a growing body of work indicating that the DMN is not simply a task-negative network (Spreng, 2012), but suggests that the DMN interacts with other brain regions engaged in task performance, specifically the striatum during reward processing in the current analysis.

**Figure 3:**
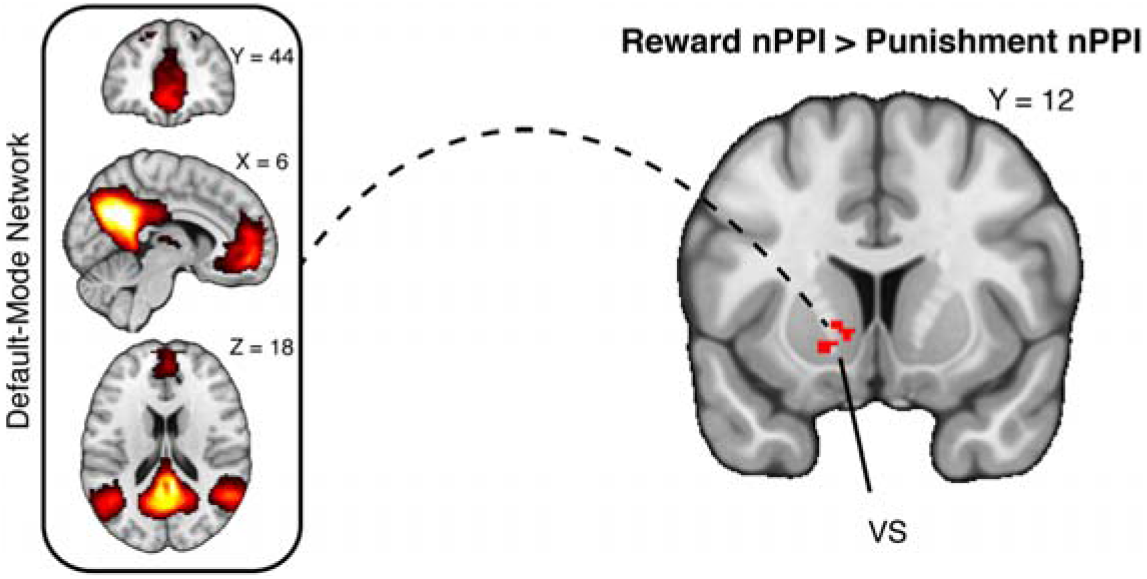
Enhanced Coupling between DMN and Ventral Striatum During Reward. We examined task-dependent changes with the DMN using a variant of psychophysiological interaction analyses. We found that an area of ventral striatum exhibited increased connectivity with the DMN during reward relative to punishment. NeuroVault https://neurovault.org/collections/10921/

Since the DMN has been shown to contribute to a spectrum of behaviors and psychological processes, we evaluated the functional significance of DMN-VS connectivity by assessing whether discounting and personality measures are significant predictors of DMN-VS connectivity. Results of the linear regression (Table 2) indicated that delayed discounting was not a significant predictor of DMN-VS connectivity. However, the constructs of openness (ß=. 138, p=.004) and agreeableness (β=-.124, p=.02) from NEO-FFI were significant predictors of DMN-VS connectivity. The NEO-FFI factor of openness has been shown to be related to sensation seeking and to reflect openness to experience, with low and high openness being characteristic of psychiatric disorders (Piedmont, Sherman, Sherman, Dy-Liacco, & Williams, 2009; Whiteside & Lynam, 2001). While there has been substantial research into the relationship of impulsivity (as measured by delayed discounting) with brain activity and connectivity (Chen, Guo, & Feng, 2017) (Hobkirk, Bell, Utevsky, Huettel, & Meade, 2019; Li, et al., 2013), investigations of the relationship between personality characteristics and brain activity/connectivity are in its early stages.

**Table 2.** Multiple regression analysis results

| Predictors | Beta | t | p |
| --- | --- | --- | --- |
| **NEO-FFI Agreeableness** | **-0.124** | **-2.404** | **0.017** |
| **NEO-FFI Openness to Experience** | **0.138** | **2.881** | **0.004** |
| NEO-FFI Conscientiousness | 0.043 | 0.803 | 0.422 |
| NEO-FFI Neuroticism | -0.045 | -0.841 | 0.401 |
| NEO-FFI Extraversion | 0.036 | 0.676 | 0.499 |
| Delay Discounting Area Under the Curve for \$200 | 0.031 | 0.48 | 0.631 |
| Delay Discounting Area Under the Curve for \$40,000 | -0.035 | -0.548 | 0.584 |
| Pos > Neg activation | 0.022 | 0.468 | 0.64 |
| Mean Relative Motion | -0.03 | -0.626 | 0.532 |

To examine whether our result of DMN-VS connectivity depends on the DMN being a continuous network, we conducted a follow up seed-based PPI analyses using representative nodes of the DMN. Specifically, we used the ventromedial PFC (VMPFC), posterior cingulate cortex (PCC), and left and right temporo-parietal junction (lTPJ and rTPJ, respectively) as seeds. These four additional PPI analyses failed to reveal reward-dependent connectivity between individual DMN seeds and the VS, indicating that the relationship between the DMN and the VS depends critically upon analyzing the DMN as a continuous network (cf. Smith et al., 2014).

We conducted additional analyses to evaluating the specificity of DMN findings by examining the connectivity between the VS and other networks. Repeated measures ANOVA on connectivity parameters from each network showed the DMN to have the strongest connectivity with the VS (F(453,1)=17.122, p < 0.0001). Post-hoc paired t-tests revealed that the DMN-VS connectivity was significantly greater than ECN-VS connectivity (t=3.3, p<.005), rFPN-VS connectivity (t=5.8, p<.0001), and lFPN-VS connectivity (t=5.8, p<.0001), pointing to the specificity of DMN-VS coupling during reward processing in comparison to other networks.

## Discussion

With decision making and processing of action-outcome associations being vital in everyday life, it is imperative to understand the involvement of brain networks in these core processes, since they are purported to play an integral part in neural organization. Here, we examined the functional relationship between the DMN and processing of rewarding outcomes in a large sample of healthy individuals. Our findings demonstrate that the receipt of reward enhances effective connectivity between the DMN and the VS. The strength of connectivity was correlated with personality characteristics of openness and agreeableness. These results advance models of DMN by demonstrating how it contributes to other brain systems and how those contributions relate to individual differences.

Our results also point to the dynamic nature of the DMN. Although many studies have shown that the DMN is deactivated during many task contexts, relatively less is known about how connectivity with other brain regions changes during these deactivations. Our results show that DMN connectivity with the VS is enhanced during receipt of reward but suppressed during the receipt of punishment. Such results build on prior studies highlight the complexity of DMN connections during tasks states (Utevsky et al., 2014; Leech et al., 2011).

Our analysis also revealed that the strength to the DMN-VS connectivity during reward processing is positively associated with the personality factor of openness and agreeableness. While we did not have an a *priori* hypothesis about the directionality of the association between connectivity and personality, this result contributes to the growing body of literature that investigates brain correlates of various personality types in both healthy (R. E. Beaty, et al., 2018; Sampaio, Soares, Coutinho, Sousa, & Gonçalves, 2014b; Wei, et al., 2014) and clinical populations (Takahashi, et al., 2013). Further, recently openness has been shown to be associated with increased functional connectivity and dopamine release from the substantia nigra (Passamonti, et al., 2014; Suridjan, et al., 2012), a subcortical brain area that is heavily involved in reward processing and projects to the striatum (Haber & Knutson, 2010).

Understanding how reward enhances connectivity between the striatum and DMN may also shine new light on prior work that has linked DMN function to disease and psychopathology at rest and during task performance (Anticevic, et al., 2012; Dobryakova, Rocca, & Filippi, 2018). Notably, depression and schizophrenia have been linked to aberrant patterns of resting-state functional connectivity between the DMN and striatum. For example, individuals with schizophrenia show reduced resting-state functional connectivity between the DMN and the striatum (X. Wang, et al., 2015). Similar effects have been observed in individuals at risk for psychosis (Hua, et al., 2019). Although altered DMN-striatal connectivity during resting state has also been observed in depression, the results are more mixed: some studies report hypoconnectivity relative to healthy controls (Bluhm, et al., 2009); while other studies report hyperconnectivity relative to healthy controls (Hwang, et al., 2016). These studies highlight the importance of DMN-striatal connectivity in disease and psychopathology, but we lack a unified theoretical framework for understanding how these aberrant forms of DMN-striatal connectivity contribute to behavior and task performance. Our results provide an important step forward by characterizing the role of reward in enhancing connectivity between the DMN and the striatum during task performance and show how this task-dependent connectivity is associated with personality variables.

Nevertheless, our results are also accompanied by limitations that merit further discussion. First, although our results may reflect a context-specific (reward receipt) modulation of effective connectivity from the DMN to the VS, such findings could also be explained by a modulation of stimulus-specific responses in the VS (K. J. Friston, et al., 1997). Specifically, responses in the DMN may enhance the striatal response to reward without a change in connectivity. Future studies may be able to disambiguate these alternative explanations using dynamic causal modeling (K. J. Friston, et al., 2003) or noninvasive brain stimulation (Polania, Nitsche, & Ruff, 2018). Second, it is not clear how our reported links between DMN-VS connectivity and personality factors would generalize to more diversified samples. Indeed, the HCP consists of healthy young adults, which may contribute to the limited range in personality scores and relatively small effect size in the brain-behavior correlations (Cremers, Wager, & Yarkoni, 2017; Yarkoni, 2009). Third, our results are limited to the narrow construct of reward consumption (D. V. Smith & Delgado, 2015). Examining other domains of reward processing—including anticipation (Knutson, Westdorp, Kaiser, & Hommer, 2000), learning (Daw, Gershman, Seymour, Dayan, & Dolan, 2011; Dobryakova & Tricomi, 2013), and valuation (Arulpragasam, Cooper, Nuutinen, & Treadway, 2018)—may reveal other patterns of DMN-VS connectivity or connectivity with other networks (Hallquist, Geier, & Luna, 2018). In a similar vein, it will be important for future work to examine how DMN-VS connectivity during reward processing is modulated by contextual factors (Dennison, Sazhin, & Smith, 2022), such as perceived control (K. S. Wang & Delgado, 2019), effort (Dobryakova, Jessup, & Tricomi, 2017), and social context (D. S. Fareri & Delgado, 2014).

Despite these caveats, our results support the conclusion that DMN plays a role in contributing to responses in the VS during the execution of a reward processing task. These results broaden our conceptualization of DMN function by going beyond simplistic activation/deactivation schemes and show how responses in other brain regions are influenced by the engagement of DMN. More efforts to understand the dynamic interplay between the DMN and other brain regions could lead to a more integrative and parsimonious view of its function (Dohmatob, et al., 2020). Such efforts could also help understand how the DMN contributes to clinical syndromes such as schizophrenia (Hu, et al., 2017), depression (Posner, et al., 2016), and substance abuse (Zhang & Volkow, 2019), potentially providing new avenues for treatment and interventions.

## Acknowledgments

This work was supported, in part, by National Institutes of Health grants R21-MH113917 (DVS), R03-DA046733 (DVS), RF1-AG067011 (DVS), R01-NS121107 (ED). We note that DVS was a Research Fellow of the Public Policy Lab at Temple University during the preparation of this manuscript (2019-2020 academic year).

## Conflict of interest statement

The authors have no conflicts to disclose.

## Notes

### Competing Interest Statement

The authors have declared no competing interest.

### Summary of Updates

streamlined introduction, added methodological details, and clarified discussion.

